# Conserved coexpression at single cell resolution across primate brains

**DOI:** 10.1101/2022.09.20.508736

**Authors:** Hamsini Suresh, Megan Crow, Nikolas Jorstad, Rebecca Hodge, Ed Lein, Alexander Dobin, Trygve Bakken, Jesse Gillis

## Abstract

Enhanced cognitive function in humans is hypothesized to result from cortical expansion and increased cellular diversity. However, the mechanisms that drive these phenotypic differences remain poorly understood, in part due to the lack of high-quality cellular resolution data in human and non-human primates. Here, we take advantage of single cell expression data from the middle temporal gyrus of five primates (human, chimp, gorilla, macaque and marmoset) to identify 57 homologous cell types and generate cell-type specific gene coexpression networks for comparative analysis. While ortholog expression patterns are generally well conserved, we find 24% of genes with extensive differences between human and non-human primates (3383/14,131), which are also associated with multiple brain disorders. To validate these observations, we perform a meta-analysis of coexpression networks across 19 animals, and find that a subset of these genes have deeply conserved coexpression across all non-human animals, and strongly divergent coexpression relationships in humans (139/3383, <1% of primate orthologs). Genes with human-specific cellular expression and coexpression networks (like *NHEJ1, GTF2H2, C2* and *BBS5*) typically evolve under relaxed selective constraints and may drive rapid evolutionary change in brain function.

**One Sentence Summary:** Cross-primate middle temporal gyrus single cell expression data reveals patterns of conservation and divergence that can be validated with population coexpression networks.

## Main Text

Cortical expansion and increased cellular diversity in the human brain following divergence from great apes are hypothesized to contribute to enhanced cognitive function (*1, 2*), but the molecular mechanisms underlying human brain evolution are not fully understood. High protein sequence conservation between humans and non-human primates suggests that cortical evolution in the human lineage is driven primarily by changes in gene expression regulation (*3, 4*). Comparative cross-species transcriptomic analyses are essential to uncover gene expression programs underlying cell identity (*5–7*), and assess the impact of their dysregulation in neuropsychiatric disease (*8, 9*). Difficulty in obtaining and preserving samples, and the poor quality of genome annotation in non-human primates have restricted the scope of most comparative studies in primates to characterizing patterns of gene regulation across a small set of species using bulk transcriptomic data from a limited number of tissues (*10–14*). Moreover, recent analyses (*15, 16*) highlight the difficulty of disentangling functional gene co-regulation confounded with coexpression due to variation in cell type abundance across tissue samples. Comparative coexpression analysis at single cell resolution has the potential to systematically trace the origin and diversity of cell types across animal evolution.

Single-cell transcriptomics has become a powerful tool to identify regional and interspecific variation in gene expression underlying the evolution of brain regions and cell types within (*17–19*) and across species (*20–23*). Comparing coexpression networks derived from single-cell profiling of matched cortical samples across different phylogenetic groups is key to identifying human-specific patterns of gene activity driving brain evolution. Importantly, aligning human and mouse samples from homologous brain regions revealed extensive divergence in gene expression of cortical cell types (*2*), and the presence of primate-specific striatal interneuron population (*24*), highlighting the importance of studying primate brains at high resolution to uncover mechanisms behind evolutionary innovations in the human lineage.

Improved genome annotations in apes and macaques (*25, 26*), and advances in single cell sequencing technologies have resulted in a steady growth in primate functional genomics studies in recent years (*7, 27–29*). The Brain Initiative Cell Census Network (BICCN) consortium has generated high quality, single nucleus transcriptomic atlases of the middle temporal gyrus (MTG) sampled across five primates spanning a ~ 45 million year evolutionary period (human, chimp, gorilla, macaque and marmoset) in a large-scale effort to explore the evolution of cellular diversity across primates at unprecedented resolution. The essence of our approach was to identify cell types shared across species and then use this common sample space to determine where and how orthologs change their expression pattern. In this study, we identified 57 homologous cell types by aligning single-nucleus MTG atlases of five primates. We observed high cross-species similarity of expression variability over 57 consensus cell types for all orthologous genes, suggesting conserved transcriptional patterning across primates. Most single-cell comparative studies have focused on differential gene expression across species to isolate species-specific changes in gene activity suggesting functional divergence. Since gene regulatory programs shape and define cell identity, we compared gene coexpression networks across species to understand how cell identity is maintained, and how it evolves. Transcriptional programs defining cell identity were highly conserved across species, with their extent of conservation following evolutionary divergence times.

Interestingly, we found 24% of 14,131 genes with conserved expression variation between human and non-human primates in a few cell classes, and divergent expression variation in other cell classes. These genes were enriched for synapse assembly and function, and nearly half of the genes showed expression divergence limited to glial cell types. To assess whether these observed changes in single cell data were robustly capturing evolutionary changes in the regulatory landscape, we used better powered bulk population coexpression networks. For each gene, we measured the reproducibility of their top coexpression partners in meta-analytic aggregate coexpression networks of 19 animals (*30*) across several thousand samples, and observed signatures of divergent regulation exclusive to the human lineage in 139 genes. Overall, we show that integrative analysis of gene expression and regulation at different levels of cellular organization is a powerful approach to distinguish evolutionarily conserved transcriptional features from uniquely human gene expression traits.

## Results

### Consensus MTG taxonomy across primates

The Brain Initiative Cell Census Network generated high resolution transcriptomic maps of the middle temporal gyrus in human, chimpanzee, gorilla, macaque and marmoset by applying single-nucleus transcriptomic (snRNA-seq) assays to samples isolated from 3 to 7 donor brains in each species (plate-based SMART-seq v4 (SSv4) for great apes, in addition to droplet-based Chromium v3 (Cv3) RNA sequencing for all primates). In total, 574,156 nuclei passed quality control, including 341,469 excitatory (glutamatergic) neurons, 158,188 inhibitory (GABAergic) neurons, and 74,499 non-neuronal cells (**Fig. 1A**).

**Fig. 1.**
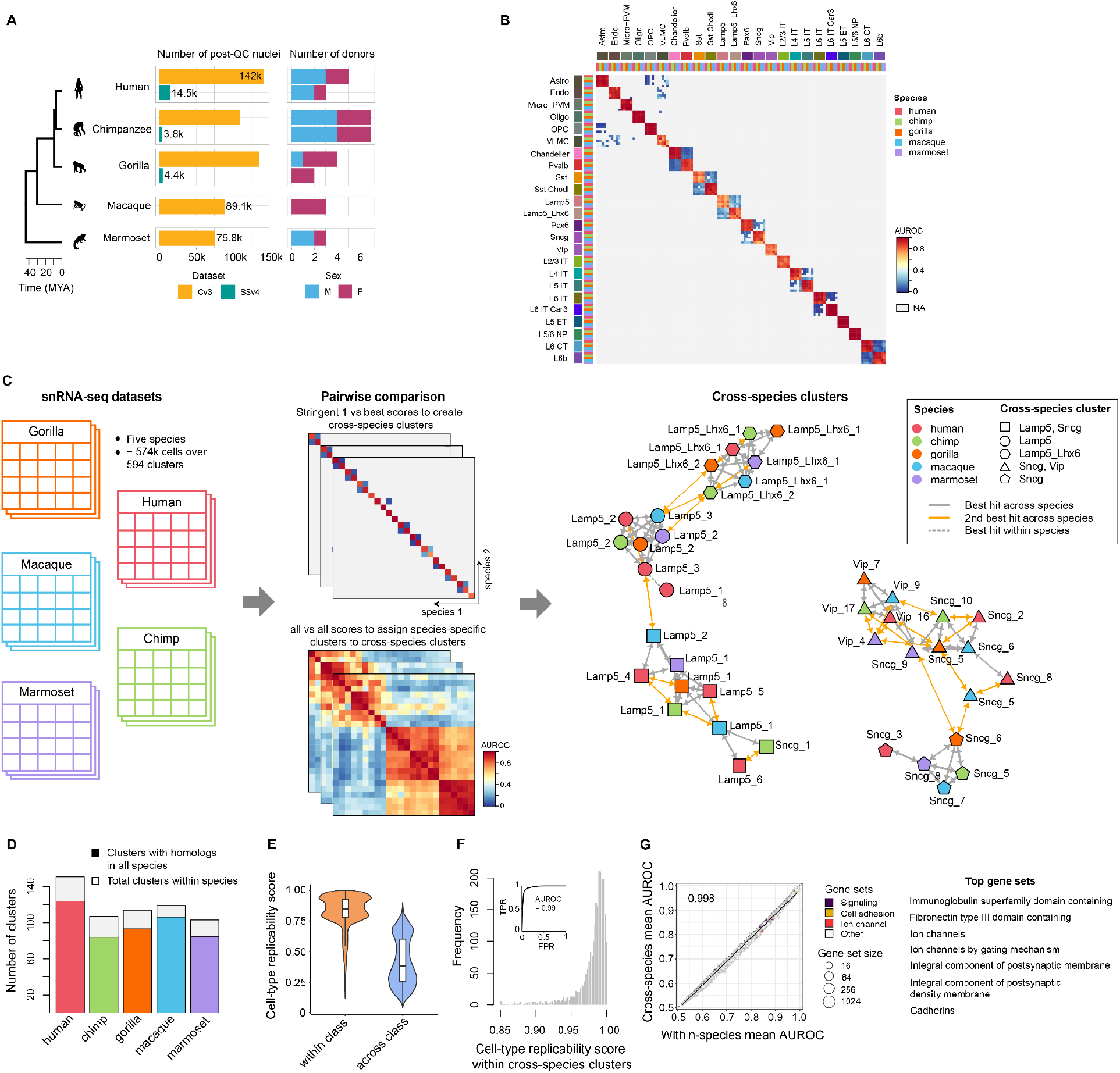
Homologous cell types across five primates. (**A**) Summary of single-nucleus transcriptomic data split by sequencing technology, number and sex of donors for each species. (**B**) Heatmap shows reproducibility of cell types across primates, with cell types labeled by species and subclass. (**C**) Schematic shows a semi-supervised MetaNeighbor framework used to define consensus transcriptomic cell types across primates. (**D**) Fraction of cell types from each species in the consensus MTG taxonomy. Cross-species clustering of cell types is validated by (**E**) comparing cell type reproducibility within and across cell classes, and (**F**) by plotting the distribution of cell type replicability scores for matched clusters across species. The ROC (Receiver-Operating Characteristic) curve in the inset indicates cluster replicability score identifies consensus cell types with high fidelity. (**G**) Scatter plot depicts the performance of 920 HGNC- and SynGO-curated gene groups in classifying consensus cell types within and across species, colored by functional category. Linear regression fit is indicated by the black line, with the slope in the upper left corner. Top highly conserved gene sets across primates are listed on the right (cell type classification performance > 0.95).

For each species, cell type annotations defined on the basis of unsupervised clustering of snRNA-seq datasets at different levels of granularity were obtained from the BICCN. To assess the replicability of cell classes and subclasses across species, we used MetaNeighbor (*31, 32*), which identifies cell types with highly similar transcriptional signatures within and across species. Cells in each species were categorized into three classes (non-neurons, excitatory and inhibitory neurons) and 24 subclasses, and were near-perfectly replicable across species, confirming that cell types have distinct transcriptomic profiles that distinguish them at broad levels of cell classification (**Fig. 1, B and E**). However, at finer resolution, the number of cell type clusters varied from 103 in marmoset to 151 in human, with multiple clusters exhibiting transcriptomic similarities within and across species (summarized in **Table S1**). We generated a comprehensive set of homologous cell types (i.e. cross-species clusters) as described below.

First, we applied MetaNeighbor to identify highly replicable clusters across species which formed the initial pool of consensus cell types. Next, we used a weighted nearest neighbor approach to assign each of the remaining unmapped clusters to the consensus cell type containing the majority of transcriptionally similar cell clusters (**Fig. 1C**; see Methods for more details). This clustering procedure allowed us to map 594 clusters in all five primates to 86 cross-species clusters, with each cross-species cluster containing one or more clusters from at least two primates. All primates shared 57 of 86 cross-species clusters (**Fig. S1**). We refer to these shared clusters as homologous cell types, and they contain more than 80% of clusters from each species (**Fig. 1D**). As expected, homologous cell types showed similar transcriptional profiles across species (**Fig. 1F**). We also functionally characterized our consensus clusters by identifying HGNC- and SynGO-curated gene groups that contributed the most to replicability (*32*). Genes related to cell adhesion and neuronal signaling were most informative of cell type identity, and showed similar classification performance when trained and tested in the same or different species (**Fig. 1G**; scores for all 920 gene groups listed in **Table S2**).

### Conserved regulatory landscape across primates

We organized the 57 homologous cell types into a hierarchical taxonomy on the basis of transcriptomic similarities (**Fig. 2A**), and observed that hierarchical relationships among cell types roughly mirrored their developmental origins. This consensus taxonomy provides an excellent opportunity to infer the extent of functional conservation between humans and non-human primates by comparing the similarity of gene expression signatures across homologous cell types.

**Fig. 2.**
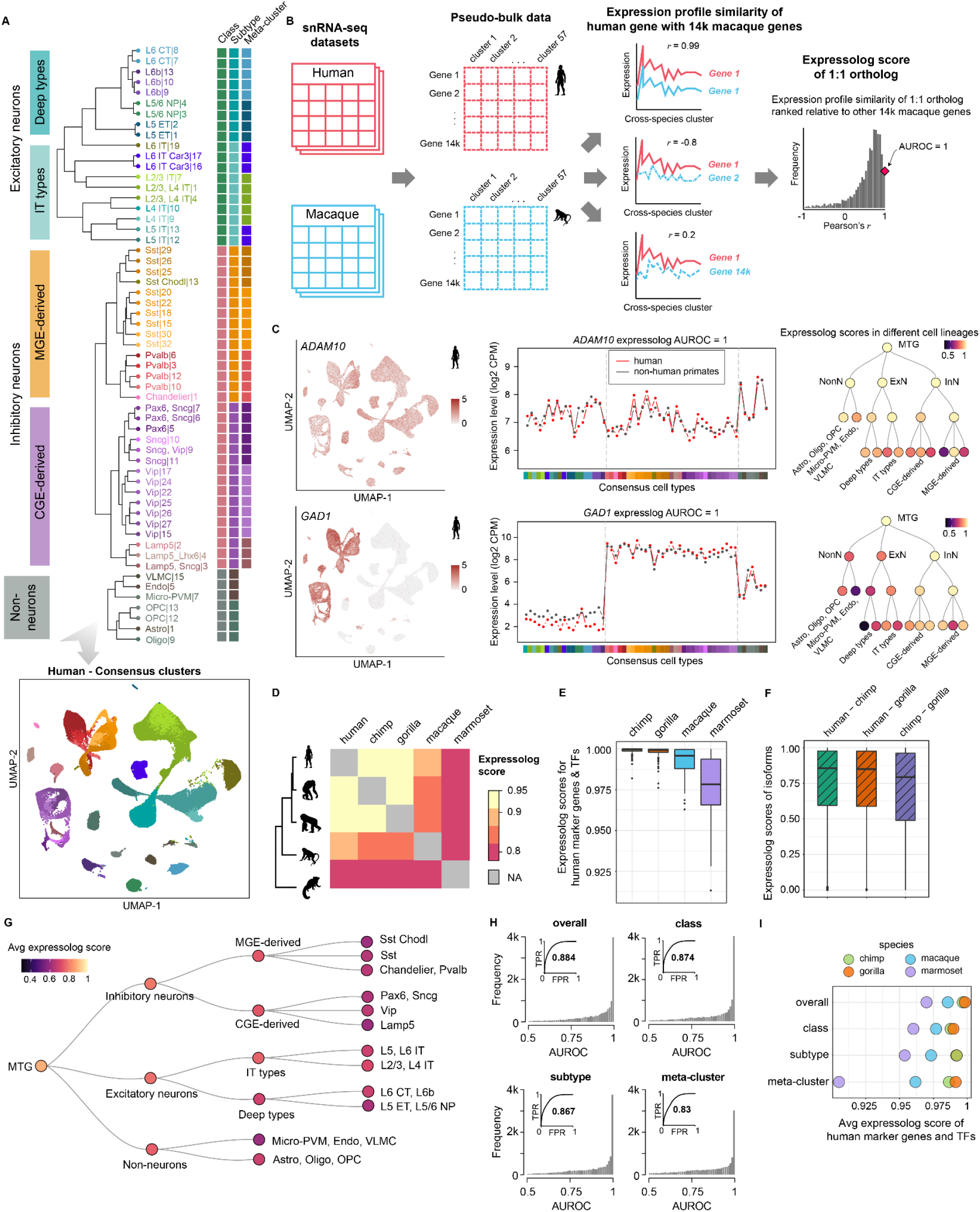
Orthologs have conserved expression profiles across primates and the extent of conservation recapitulates known phylogeny. (**A**) (top) Dendrogram of 57 consensus cell types defined by their transcriptomic similarity, annotated with corresponding cell class, subtype and meta-cluster; (bottom) UMAP plot of single nuclei from human MTG integrated across donors and snRNA-seq technologies, and colored by consensus cluster. (**B**) Schematic represents the method to calculate expression profile similarity of 1:1 orthologs for a pair of species. (**C**) Examples of genes with constitutive (*ADAM10*) and cell-type specific expression (*GAD1*) in the human MTG data (color indicates expression level in the UMAP plots). Both genes have near-identical expression profiles between human and non-human primates (expressolog score = 1 in both cases). Expressolog scores computed across cell types within each subgroup reveals transcriptomic divergence in specific meta-clusters (*Pvalb* cell subclass for *ADAM10*, and L5/6 excitatory neurons and vascular cells for *GAD1*). (**D**) Heatmap shows the distribution of expressolog scores for 14k orthologs across primates. (**E**) Boxplots indicate that lineage-specific genes like marker genes and transcription factors have conserved expression profiles across primates suggesting conserved transcriptional programs shape cell identity across species. (**F**) Expressolog scores suggest that individual isoforms also exhibit similar expression profiles across great apes. (**G**) Tree shows cell type meta-clusters organized into broad cell lineages color-coded by average expressolog scores between humans and non-human primates. (**H**) Histograms show the distribution of average expressolog scores between humans and non-human primates stratified by cell type hierarchy. Insets - expression profile similarity predicts orthologs across primates at multiple scales, as seen by ROC curves stratified by cell type hierarchy. (**I**) Marker genes and transcription factors remain expressologs at different cell type resolutions, with average expressolog score decreasing with phylogenetic divergence and cell type homogeneity.

Adopting the language of Patel et al (*33*), given a query gene from one species, the homologous gene in the target species with the most similar expression variability across a set of matched tissues is referred to as its “expressolog”. While this method has been employed to select functionally similar orthologs from homologous gene clusters, we apply it here to evaluate the similarity of expression profiles of 1:1 orthologs compared to that of random gene pairs. We obtained a list of 14,131 human genes with one-to-one (1:1) orthologs in all non-human primates from OrthoDB (*34*). For each pair of species, we calculated the expression profile similarity for all pairs of genes by correlating the mean normalized expression levels across 57 homologous cell types. We then defined the rank-standardized expression profile similarity of 1:1 orthologs relative to all other genes as the “expressolog score” (see **Fig. 2B** for schematic representation, and Methods for details on calculation). In essence, this measures whether orthologs show similar expression profiles across cells. This score is represented as an AUROC (Area Under the Receiver-Operating characteristic Curve) with a score of 1 signifying specific and highly similar expression variation across the species pair, 0.5 indicating dissimilar/uncorrelated expression variation and 0 indicating significant extreme expression profile divergence in one or both species.

The expressolog score for each gene measures the specificity with which transcriptional signatures across shared cell types can be used to detect its 1:1 orthologs across species. Intuitively, an expressolog score of 0.99 for a gene indicates that its ortholog is in the top 1% of all genes in terms of expression profile similarity. Since genes with shared functions often display similar expression profiles, we use expressolog scores computed over 57 matched cell types as a measure of gene functional conservation across species. We find that orthologous genes show highly similar patterns of expression variation across cell types and are highly conserved across the phylogeny. Two such examples are shown in **Fig. 2C**: *ADAM10*, which is constitutively expressed in the primate MTG, and *GAD1*, which is expressed exclusively in inhibitory neurons. Both genes exhibit perfectly matched expression profiles across human and non-human primates, corresponding to an expressolog score of 1 in each case. Remarkably, both *ADAM10* and *GAD1* display conserved patterns of expression variation even across more homogeneous cell types, suggesting that the genes are deeply conserved across species.

Early microarray-based comparative studies noted that the divergence in gene activity in the same tissue between species reflects their evolutionary relationships (*12, 35*). Likewise, average expressolog score (or cross-species coexpression) correlates with evolutionary distances between human and non-human primates (**Fig. 2D**; refer **Table S3** for the full list of scores). Expressolog scores correctly classify orthologs with performance ranging from 0.93 for humans with great apes, to 0.8 for humans with marmoset. Marker genes and transcription factors (TFs) also show high functional conservation across species, suggesting a highly conserved molecular landscape of the MTG region across primates (**Fig. 2E**, see Methods for details on marker and TF selection).

Alternative splicing is known to increase transcriptomic diversity in primates (*36, 37*), but the functional conservation of individual isoforms is yet to be fully characterized. Do isoforms have reproducible transcriptional signatures across primates? To address this, we used SSv4 data with full transcript coverage in 28 cell types in human, chimp and gorilla to explore patterns of isoform usage across great apes. In general, isoforms showed similar expression profiles across species (**Fig. 2F**). For each gene with multiple isoforms in a pair of species, we calculated the expressolog scores for all isoform pairs to measure the ability of each isoform to correctly predict itself across species. Overall performance for this task was slightly better than that expected by chance, suggesting similar but not specific transcriptional patterning of isoforms across species (AUROC = 0.56). Indeed, consistent with previous observations (*37*), we also observed extensive isoform switching across apes which could explain the weak expression specificity of isoforms across species.

Since cell classes are transcriptionally distinct, genes typically have highly variable expression across cell classes which could drive their strong cross-species coexpression. Does the expression profile similarity persist at finer cell type resolution? Based on their transcriptomic and spatial similarity, we recursively subdivide the consensus cell types in the three classes into 6 subtypes (2 from each major class) and 10 meta-clusters (3 from each inhibitory subtype and 2 from each excitatory subtype), and evaluate the expression profile similarity of orthologs within these subsets of the cell type hierarchy (**Fig. 2A**). We observe consistent cross-species coexpression of orthologs at multiple scales (AUROC = 0.83 within meta-clusters and 0.87 within classes) suggesting a core co-regulatory network shaping cell identity across different scales of cellular organization (**Fig. 2, G and H**; see Methods and **Fig. S2** for more details).

Marker genes and transcription factors known for cell type-specific expression also show significant covariation with cell types in other cell lineages (AUROC = 0.96 within meta-clusters and 0.98 within classes; **Fig. 2I**), consistent with similar analyses of the BICCN mouse primary motor cortex data (*38*). We hypothesize that tightly coordinated, differential regulation of functionally conserved genes creates graded, continuous variations in expression levels across cell types, resulting in persistent cross-species coexpression at different scales of cellular organization (*39*). As an example, we show that the inhibitory neuron marker *GAD1* exhibits graded expression differences across cell types, yet remains strongly correlated across species within cell types (**Fig. S3**). Importantly, genes defining cell identity show differences in expression profile mirroring primate phylogenetic relationships at all levels of granularity.

### Coexpression neighborhood similarity suggests conserved gene activity across species

So far, we have used gene expression profile similarity as a measure of functional conservation across species. Since gene coexpression reflects shared function or regulation, a complementary approach to assess functional conservation of orthologs is to quantify the similarity of their broader coexpression neighborhoods across species (*40–43*). We built gene coexpression networks for each of the five primates by aggregating individual coexpression networks built from pseudo-bulk samples of 57 cell types (see Methods for more details). We then compute (i) the connectivity of marker gene modules in each network using a neighbor-voting method (*44*), and (ii) coexpression neighborhood conservation of 1:1 orthologs across each pair of networks (*43*), as discussed below (see **Fig. 3A** for schematic representation).

**Fig. 3.**
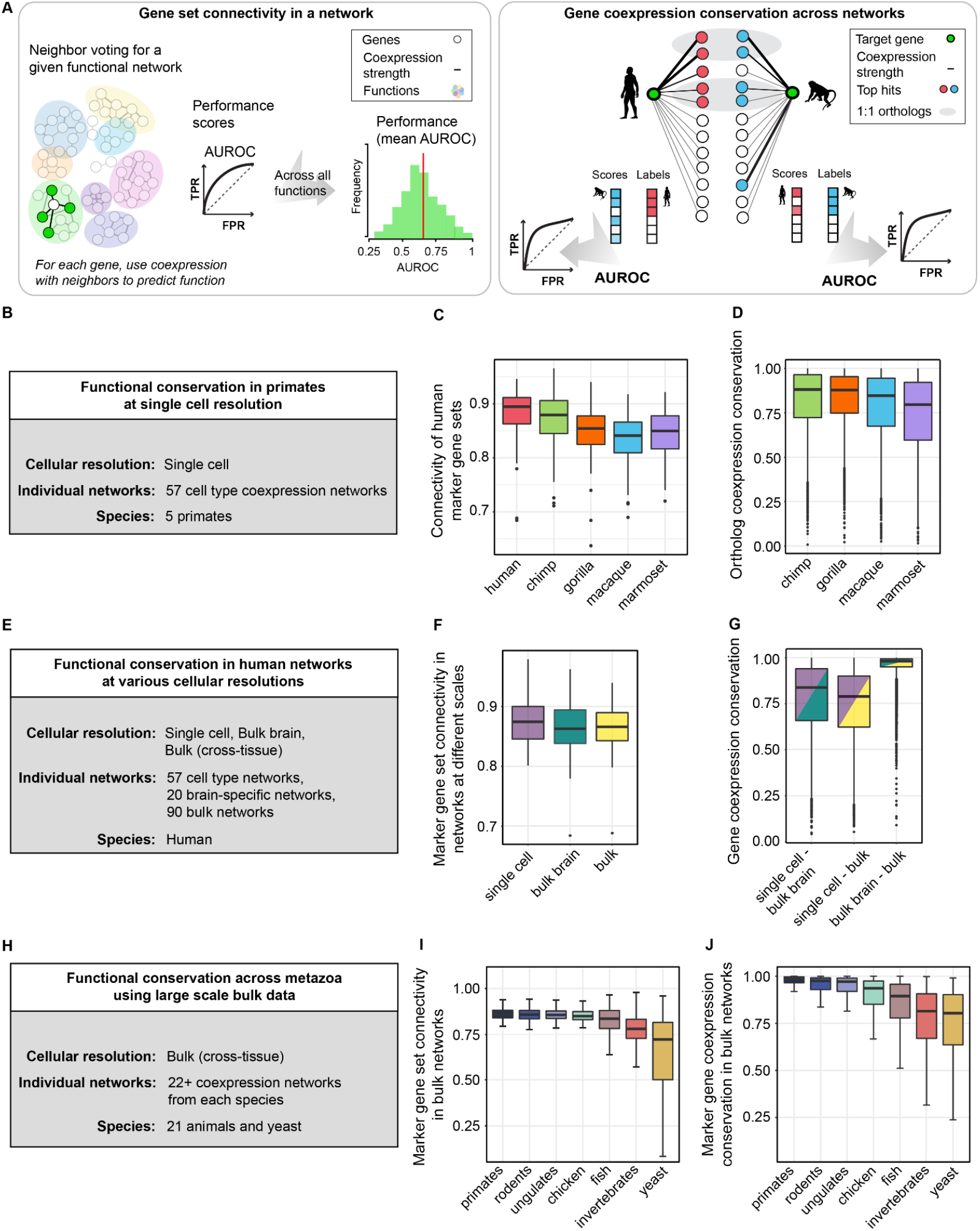
Coexpression neighborhood similarity highlights conserved gene regulatory landscape across metazoa. (**A**) Functional conservation is assessed by (i) a neighbor-voting method used to assess the modularity of gene sets in our aggregate coexpression networks (left panel), and (ii) conservation of coexpression neighborhoods for 1:1 orthologs of a species pair (right panel). (**B**) Aggregate coexpression networks at single cell resolution are built by aggregating 57 cell type-specific coexpression networks in each primate. (**C**) Boxplot indicates that marker genes sets form highly connected modules in primate single cell coexpression networks. (**D**) Boxplots show the distribution of coexpression conservation for 4500 highly variable genes calculated between human and non-human primates. (**E**) Brain-specific and cross-tissue coexpression networks generated from 20 RNA-seq datasets in Gemma and 90 datasets in SRA respectively, to assess the conservation of coexpression relationships at various levels of cell type heterogeneity. Boxplots show (**F**) conserved topology of networks across different scales of cellular organization, and (**G**) strong coexpression conservation between “compositional” (bulk) and “co-regulatory” (single cell) networks. (**H**) Coexpression networks generated from 22 or more RNA-seq datasets from SRA for each of the 22 species and downloaded from CoCoCoNet. (**I**) Boxplots indicate that marker genes defining consensus cell types show high modularity in networks with high compositional heterogeneity, and across evolutionarily distant species. (**J**) Boxplots show mean coexpression conservation for marker genes between human and other species, grouped by their divergence time. Coexpression conservation is negatively correlated with phylogenetic distance (rho=-0.65, p<10^-16^).

Since these networks are built from relatively homogeneous cell populations and capture expression variation only within cell types (**Fig. 3B**), we were surprised to find that genes defined as markers for heterogeneous cell types (class, subtype or meta-cluster) formed highly connected modules in all species networks (**Fig. 3C**). This confirms the presence of core co-regulatory networks even across highly refined cell types as suggested by previous studies examining cell type-specific coexpression networks in mouse (*38*) and fly brains (*45*).

To measure the similarity of gene coexpression neighborhoods between human and non-human primates, we subset gene coexpression networks to 4500 highly variable 1:1 orthologs, and calculate a “coexpression conservation” score, which is a measure of gene neighborhood replicability across the species pair (see right panel of **Fig. 3A** for schematic representation, and Methods for further details). We observe that gene coexpression neighborhoods are highly conserved across primates, revealing a highly conserved cellular architecture of the MTG region across primates. Similar to expression profile similarity, we find that coexpression neighborhood similarity also correlates with primate phylogeny (**Fig. 3D**).

Are gene coexpression relationships replicable across networks built at different levels of cell type heterogeneity? We can now compare coexpression networks from snRNA-seq datasets with networks derived from whole brain or cross-tissue samples in humans to distinguish coexpression due to shared co-regulation from coexpression driven by cell type composition. We generated a meta-analytic human brain coexpression network by aggregating datasets of human bulk brain data sourced from the Gemma database ((*46*), **Fig. 3E**). At coarser resolution, we also obtained a high confidence human gene coexpression network from CoCoCoNet (*30*) created by meta-analysis of publicly available bulk RNA-seq datasets.

We calculated the modularity of marker gene sets in the three networks (single-nucleus MTG, brain-specific bulk, whole bulk networks) that capture cell type signals across a wide spectrum, and observed high modularity in all three networks suggesting shared topology across networks reflecting both compositional and co-regulatory gene-gene relationships (**Fig. 3F**). Coexpression neighborhood conservation calculated pairwise between the three aggregate coexpression networks revealed functional conservation of genes at all scales (**Fig. 3G**). The high degree of consistency between single-nucleus and bulk networks reaffirms a model of multiscale coexpression in the brain (*38*).

While the neocortex is a feature specific to mammals, its basic components may have evolved prior to mammalian evolution and undergone extensive reorganization in different phylogenetic classes (*47, 48*). We use coexpression conservation of functionally relevant genes to test for signs of conserved molecular identity across species. Ideally, we would like to test this idea through meta-analysis of large scale brain-specific transcriptomic data from multiple species, but such data is only available for select model species. Previously, we showed that our gold standard human gene coexpression network (assembled from datasets sampling multiple bulk tissues) is topologically and functionally similar to our meta-analytic brain-specific human network, capturing the key regulatory features shared by both. Based on this observation, we tested the conservation of neuronal and non-neuronal marker genes using bulk coexpression networks of humans and 21 other species available on CoCoCoNet (**Fig. 3H**; networks derived from 54,668 samples over 22 species as reported in **Table S6**). As expected, marker gene sets form densely connected modules in all animals, indicating ancient, conserved regulatory features across metazoa ((*5, 49*), **Fig. 3I**). We observe consistently high coexpression conservation scores of 1:1 orthologs even in phylogenetically distant species like fruit fly and roundworm (**Fig. 3J**), suggesting that the orthologs are present in the same functional modules in all species. The presence of conserved gene modules in distant species which lack homologous tissues (i.e. neocortex) indicates extensive repurposing of transcriptional programs over metazoan evolution.

Overall, our comparative coexpression analysis at different scales and across divergent species provides evidence for widespread functional conservation of genes across metazoa. For example, the inhibitory neuron marker *GAD1* is differentially expressed across cell classes (high expression in inhibitory neurons and low expression in excitatory neurons), and consequently exhibits strong coexpression across homologous cell types in primates (**Fig. S3**). However, comparative coexpression analysis stratified by cell class reveals persistent coexpression of *GAD1* within cell types of unexpected lineage (i.e. excitatory neurons), indicating the presence of shared regulatory programs across cell types. We also observe strong coexpression conservation of *GAD1* even in species without inhibitory neurons (like yeast), validating our hypothesis that conserved gene programs are reused across cell type and species phylogenies during evolution.

### Integrative analyses with large scale bulk RNA-seq data identifies genes with human-specific regulatory divergence

Genetic variation within species is known to drive regulatory and phenotypic variation across species (*12, 50*). Under a neutral model of evolution, we expect a similar constraint of evolutionary drift to apply to gene sequences and expression levels within and across species. The evolutionary trajectory of many genes follows this principle as evidenced by highly conserved expression and coexpression profiles across large evolutionary timescales. However, a few outlier genes can have expression changes due to positive selection on specific regulatory variants, lower mutational constraint, or due to environmental differences across species (*51*).

The single-nucleus MTG atlases of five primates are an excellent resource to examine gene expression differences at cellular resolution, whereas our meta-analytic bulk coexpression networks across phylogenetically diverse species can be exploited to robustly infer patterns of species-specific regulatory divergence. Here, we propose that an integrative analysis of high resolution single-nucleus and well-powered bulk transcriptomic data can combine the specificity of expression across cell types with the similarity of coexpression neighborhood across divergent species to detect human-specific regulatory variation in an evolutionary context.

Our workflow to identify genes with potential human-specific coexpression patterns is illustrated in **Fig. 4A**. Briefly, we select genes with low expressolog scores (AUROC < 0.55) within one or more classes between human and non-human primates. These 3,383 genes exhibit diverged expression profiles either in humans, non-human primates, or across all primates. To shortlist the genes with potential human-specific expression divergence, we then examine their coexpression conservation scores across 19 animals (humans and 18 other animals with >60% human orthologs), and only select genes showing significantly lower conservation between human and other animals, compared to all other pairs of animals. We identify 139 genes with concordant human-specific functional divergence in single cell and bulk transcriptomic data, a very small fraction of all genes analyzed, consistent with an evolutionarily conserved regulatory landscape across species. Genes exhibiting species-specific functional divergence between humans and other primates are the exception in our analysis, not the rule.

**Fig. 4.**
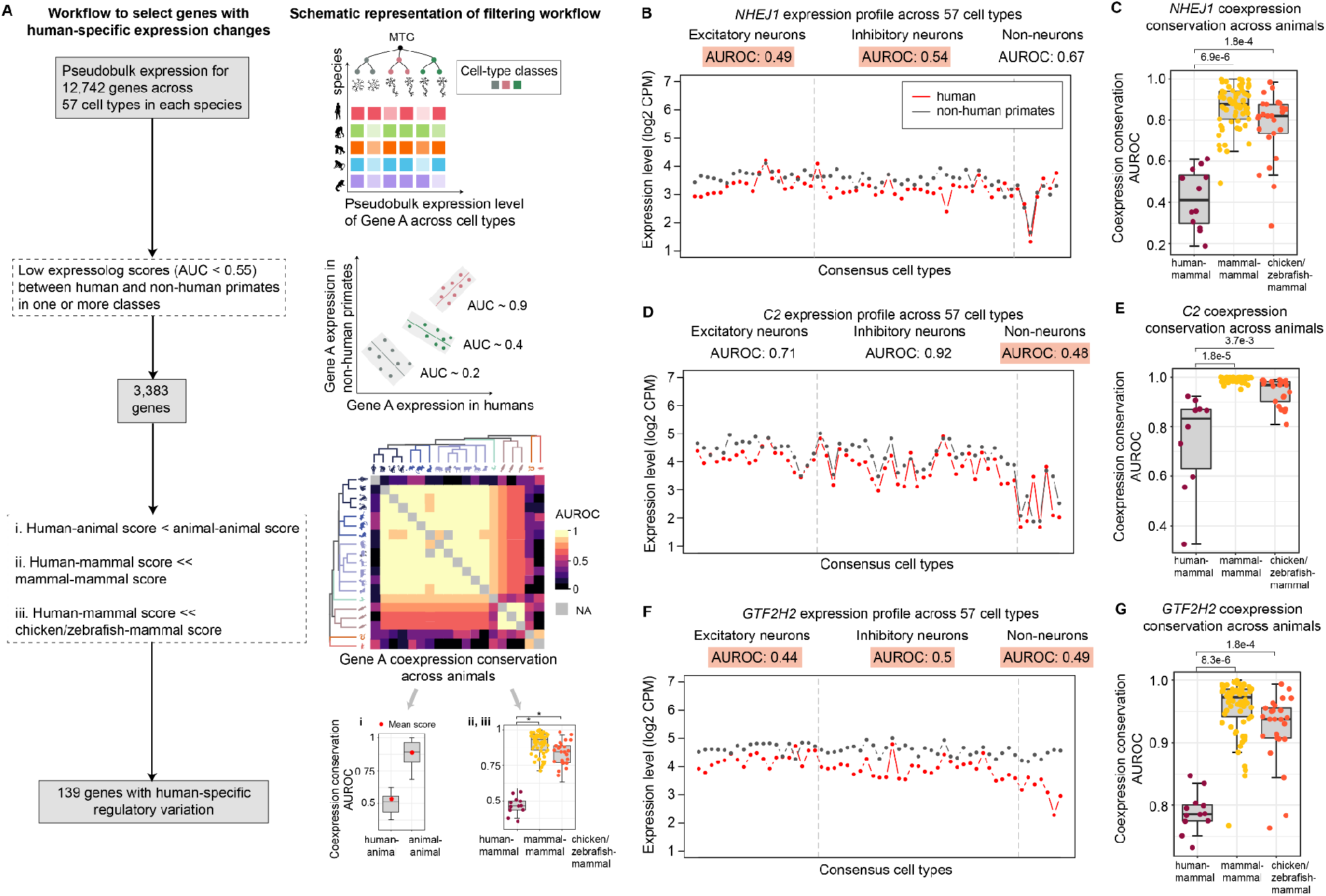
Integrative analysis of single-nucleus and bulk transcriptomic data can detect genes with human-specific regulatory divergence. (**A**) Schematic representation and workflow of our approach to identify genes with human-specific regulatory changes. We show three examples of genes displaying human-specific differential coexpression: (**B, C**) *NHEJ1*, (**D, E**) *C2*, and (**F, G**) *GTF2H2*. (**B, D, F**) Plots compare the expression profile of each gene of interest in humans with the average expression profile of the ortholog in non-human primates. Expressolog scores within each cell class are listed above the plot, and scores < 0.55 are highlighted in orange. (**C, E, G**) Boxplots show coexpression conservation for orthologs between human and non-human mammals (points colored in maroon), between pairs of non-human mammals (yellow), and between non-human mammals and other vertebrates (chicken and zebrafish; orange).

Among 3,383 genes with diverged expression profiles in one or more cell classes between human and non-human primates, 98% of genes showed expression divergence in only one cell class, with nearly half of the genes exhibiting differential coexpression across non-neuronal cell types. The 3,383 genes are more likely to be associated with cortex-specific significant expression quantitative trait loci (eQTLs; Wilcoxon *P* < 0.003) which could underlie gene expression changes in humans. Compared to all expressed genes, genes with diverged expression in one or more classes were enriched for intracellular signal transduction, synapse organization and function (Fisher’s exact test, adjusted *P* < 0.02), and significantly associated with various brain disorders including intellectual disability, microcephaly, epilepsy, autism spectrum disorders (adjusted *P* < 0.001). We detected 139 genes with putative novel regulatory relationships in humans, and a majority of these genes (68%) were diverged in a single cell class with roughly equal number of genes diverged in each of the three classes, GABAergic, glutamatergic and non-neuronal cells. The 139 human genes were also significantly associated with intellectual disability and blindness.

We visualize the expression variation over homologous cell types and cross-species coexpression conservation for three candidate genes showing human-specific deviation in expression profile in neurons (*NHEJ1*), non-neurons (*C2*), and in all cell classes (*GTF2H2*). Differences in gene expression profiles between human and non-human primates for these genes are shown in the left panels in **Fig. 4** (**B, D and F**). The boxplots on the right (**Fig. 4, C, E and G**) show the corresponding distributions of coexpression conservation between humans with non-human mammals, within non-human mammals, and between non-human mammals and other model vertebrates (chicken and zebrafish). This broader species analysis confirms differential coexpression in humans, validating the human-specific expression variation observed in single nucleus transcriptomic data.

*NHEJ1* is a DNA repair gene known to be under positive selection exclusively in the human lineage (*52*). An independent study that compiled a comprehensive list of human accelerated regions (HARs) in the genome (*53*) also identified a HAR overlapping this gene (HARsv2_1598), suggesting accelerated evolution of its coding sequence drives regulatory divergence specific to the human lineage.

Given that *cis*-regulatory variation contributes to interspecific expression divergence (*50*), association of significant expression quantitative trait loci (eQTL) with *GTF2H2* and *C2* could explain their expression variability across species. *GTF2H2* is a transcription factor gene with high inter-individual variability due to several cortex-specific eQTLs as seen in the Genotype-Tissue Expression Project (v8; (54)). *C2* is an immune-related gene involved in interferon signaling and has microglia-specific expression in the human central nervous system (*55*). *C2* is known to mediate interactions between microglia and neurons, and its downregulation in microglia is associated with aging (*56*). Since *C2* has similar expression levels in microglia and neurons, regulatory changes in the human lineage could underlie the divergent pattern of *C2* expression in non-neurons. These examples suggest the power of integrative analysis to uncover both patterns and mechanisms of human-specific expression variation, and measure the functional impact in a broad phylogenetic context.

Finally, we sought to assess genic properties of the 139 human genes that could be associated with human-specific functional divergence. Consistent with previous research (*57*), we found that the 139 genes were younger (Wilcoxon *P* < 0.006), shorter in length (Wilcoxon *P* < 3.1 x 10^-16^), had higher GC content (Wilcoxon *P* < 2.4 x 10^-5^) and displayed more cell type-specific expression (Wilcoxon *P* < 0.0003) compared to the other 12,603 functionally conserved genes. Divergent genes had marginally lower sequence similarity across primates compared to conserved genes (Wilcoxon *P* < 0.01), coupled with higher sequence evolution rates in the human lineage (Wilcoxon *P* < 0.01). Despite not being more likely to be associated with significant cortical eQTLs, divergent genes showed relatively higher tolerance to inactivation (i.e. higher LOEUF scores; Wilcoxon *P* < 8.4 x 10^-8^). These observations suggest that the divergent genes predominantly evolve under relatively mild evolutionary constraints, with a handful of genes acquiring new regulatory features (like HARs) under positive selection.

### Evidence for human-specific regulatory rewiring in the ciliopathy gene BBS5

Another gene with potential differential expression regulation specific to the human lineage is *BBS5* (Bardet-Biedl Syndrome 5). Bardet-Biedl Syndrome is a ciliopathic disorder with heterogeneous phenotypes across the human population, including retinal dystrophy, polydactyly, mental retardation, hypogonadism, obesity and kidney dysfunction, and *BBS5* is one of a family of 21 *BBS* genes linked to this disorder (*58*). *BBS5* is expressed in ciliated cells and secretes a protein which is a part of the octameric protein complex (BBSome) required for ciliary membrane biogenesis (*59*), and defects in cilium assembly or function form the basis for human ciliopathies.

Single nucleus profiling reveals significant differences in *BBS5* expression profiles between human and non-human primates, but highly similar profiles between non-human primates. We observe human-specific expression differences not just across 57 homologous cell types (heatmap in **Fig. 5A**), but also within excitatory neurons and non-neuronal cell types (boxplots in **Fig. 5A**).

**Fig. 5.**
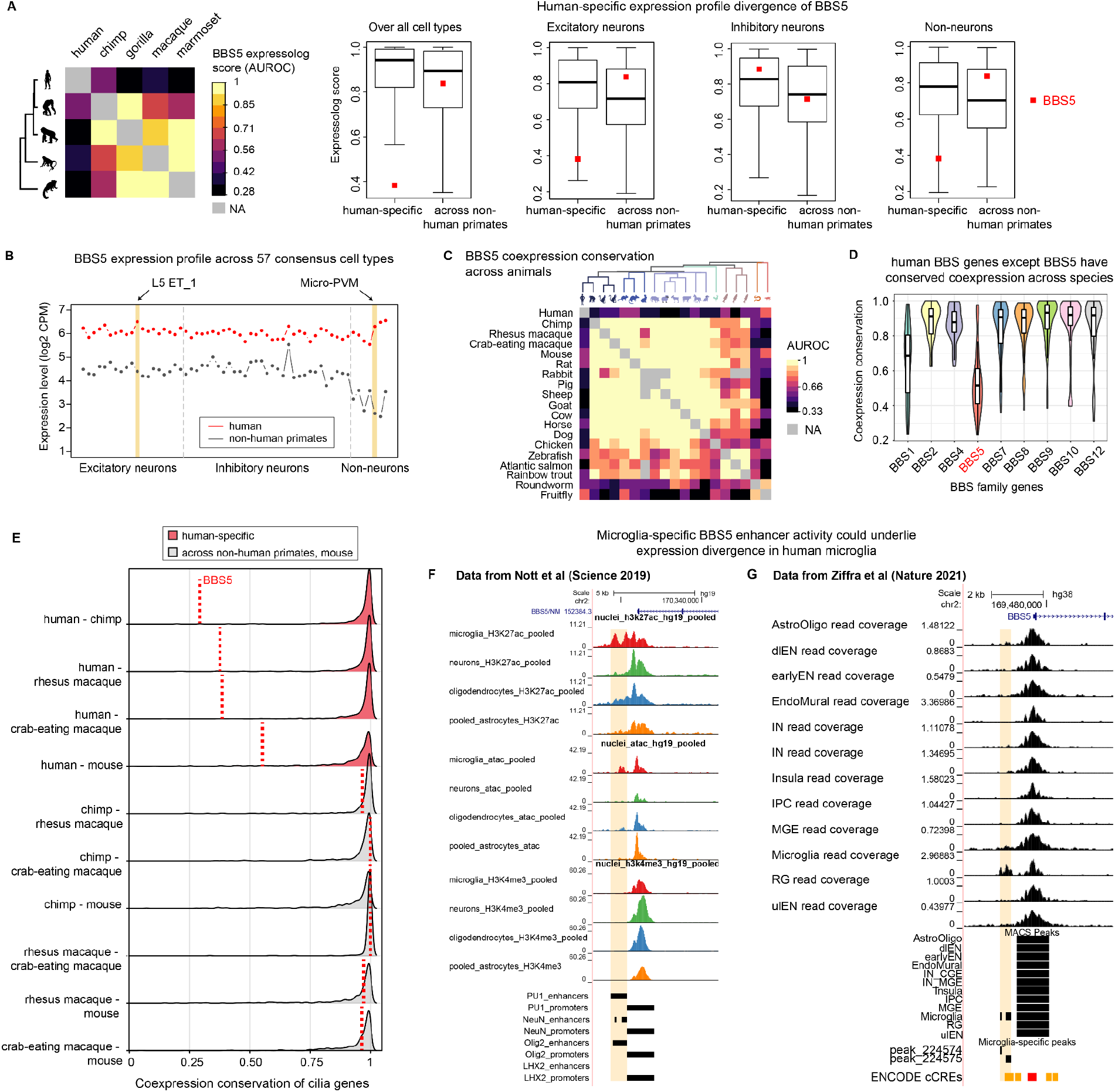
Integrative analyses with transcriptomic and epigenomic data validates regulatory rewiring of BBS5 with human-specific expression profile divergence. (**A**) (left) Heatmap shows BBS5 expression profile similarity for each primate pair, and (right) Boxplots show the expression profile similarity distribution for all orthologs both across and within cell classes, with BBS5 marked in red. Together, the plots show that BBS5 has high expression profile similarity between non-human primates, but low similarity between humans and non-human primates. (**B**) Plot shows the expression profile of BBS5 in humans with the average expression profile of its ortholog in non-human primates. BBS5 shows human-specific expression gain in neuronal (layer 5 ET neurons) and non-neuronal (microglia) cell types. (**C**) Heatmap shows high coexpression conservation specificity of BBS5 orthologs in all species pairs, excluding humans. Note that the lower scores seen for fish and invertebrates with non-human mammals is consistent with their large evolutionary distances. (**D**) Boxplot shows the distribution of cross-species coexpression conservation for 9 genes in the BBS family, with BBS5 having the least average score. (**E**) Distributions of coexpression conservation of genes involved in cilium organization and assembly across primates and mouse indicate their highly conserved function across species, with the exception of BBS5, which is conserved across non-human primates and mouse, but diverged only in humans. Single cell epigenomic profiling of broad cortical cell types in the (**F**) adult (61), and (**G**) developing human brain (62) suggests microglia-specific activity of putative BBS5 enhancer. While (**F**) also shows oligodendrocyte-specific accessibility for the enhancer, this is not replicated in (**G**).

Given that *BBS5* protein is highly conserved across primates (protein sequence similarity of 99.7% in the two great apes; data from Ensembl v107), we hypothesized that expression changes in specific cell types might be driving human-specific divergence. Indeed, we observe human-specific up-regulation of *BBS5* specifically in one layer 5 excitatory neuron cell type and in microglia (**Fig. 5B**). We also observed differential coexpression conservation between humans and other animals in our bulk networks (**Fig. 5C**). Since other *BBS* genes also exhibited distinct expression profiles in humans (see **Fig. S4** for expression divergence in human *BBS1* and *BBS10*), we tested whether *BBS* genes in general showed human-specific regulatory rewiring. Genes related to BBSome formation have been reported to be present across metazoa (*60*). Consistently, we also predict that genes involved in the formation or maintenance of the BBSome complex have conserved function across primates and rodents (**Fig. 5D**), with the exception of *BBS5*.

Cilia have ancient, evolutionarily conserved roles in embryonic development and limb patterning, and are noted as one of the cellular innovations resulting in the emergence of multicellular organisms, but are also known for their structural diversity across species (*63*). Does the structural diversity translate to network connectivity changes across species? We selected a list of 836 genes involved in cilia morphogenesis and assembly (genes from GO terms *GO:0005929, GO:0044782, GO:0060271, GO:0036064*), and compared their coexpression conservation across five species networks (human, chimp, crab-eating macaque, and two popular disease models - rhesus macaque and mouse). Cilia genes were broadly conserved across species, as expected by their evolutionarily preserved function, but human *BBS5* deviated from this trend (**Fig. 5E**), which suggests both species- and gene-specific regulatory changes. Mouse and zebrafish *BBS* mutant models are known to capture only some but not all phenotypes observed in the human disorder (*64, 65*). While Bardet-Biedl Syndrome is characterized as a genetic disorder, our results suggest that significantly low coexpression conservation of *BBS5* between humans and other animals could explain the lack of full phenotypic translation between humans and model organisms like mouse and zebrafish. As macaque models become more common to study this ciliopathy, human-specific regulatory divergence could also be related to adverse phenotypes across different tissues in humans and macaques. Further experiments are required to assess the phenotypic impact of this differential coexpression, which could in turn predict the likelihood of translational success for this gene.

Finally, we investigated the role of *cis*-regulatory elements in driving expression variability across human cortical cell types. Candidate *cis*-regulatory elements of *BBS5* (cCREs from ENCODE project) show significant sequence divergence across vertebrates, but are conserved across primates suggesting their primate-specific evolution (**Fig. S5**). Therefore, we switched our focus to search for epigenetic variability across cell types that could be associated with expression variability. Single nucleus ATAC-seq and ChIP-seq profiling of broad cortical cell types in the adult human brain ((*61*), **Fig. 5F**) show both microglia- and oligodendrocyte-specific enhancer activity in BBS5. Single cell ATAC-seq profiling of the developing human brain ((*62*), **Fig. 5G)** only shows microglia-specific enhancer activity in *BBS5*, strongly suggesting a potential mechanism for expression up-regulation in human microglia. Since new phenotypic effects of mutations in *BBS* genes continue to be reported in the literature, we suggest that microglia-specific transcriptional and epigenetic patterns could be implicated in a novel neurodegenerative disease phenotype.

## Discussion

Single nucleus transcriptomic profiling of the middle temporal gyrus in humans and four non-human primates provides an unprecedented opportunity to determine the core transcriptional features underlying conserved cell identity across primates and isolate human-specific transcriptional features related to cellular diversity and trait evolution in the human lineage. In this study, we use single nucleus expression data from the MTG of five primates (human, chimp, gorilla, macaque and marmoset) to generate a consensus transcriptomic classification of cell types, which serves as the basis for comparative analysis of gene expression across primates. Expression profile similarity of 14,131 orthologs between human and non-human primates confirmed the functional conservation of orthologs across primates (mean expressolog AUROC = 0.88; 47% of genes highly conserved with AUROC > 0.95). We also calculate the similarity of gene coexpression networks across matched cell types to assess the extent of conservation of transcriptional programs across species. Cross-species coexpression similarity reveals conserved modules across primates, with the average extent of conservation recapitulating phylogenetic distances.

Comparative analyses of large-scale meta-analytic aggregate coexpression networks (derived from 49,796 RNA-seq samples spanning 19 animals) reveal broadly conserved transcriptomic signatures across metazoa (mean coexpression conservation of 14,131 genes between humans and other animals = 0.89). Highly conserved orthologs identified from primate-specific single nucleus transcriptomic data (expressolog AUROC > 0.95) are also conserved across larger evolutionary timescales (mean coexpression conservation between human and other animals = 0.91). Interestingly, we observe that transcriptional programs defining cell identity (i.e., marker genes and lineage-associated TFs) also exhibit persistent expression variation within cell types defined at finer resolutions in primates, and show high coexpression conservation even in evolutionarily distant species lacking homologous cell types. These observations point towards: (i) the presence of conserved transcriptional modules across species and cell type phylogenies, and (ii) graded, continuous variation in expression within and across cell types defines cell identity.

One of the main goals of comparative analysis using high resolution, multi-omic profiling of matched brain regions across species is to develop methods for robust inference of genes with human-specific regulatory divergence underlying phenotypic novelty. Given that genes typically have matched expression profiles across primates, differences in co-variation likely reflect functional divergence between species. Therefore, we use cross-species coexpression between human and non-human primates within the three broad classes to identify genes with differential regulation in humans. We find 3,383 genes with divergent transcriptional patterning (expressolog score < 0.55) across one or more classes in humans relative to non-human primates. Genes diverged in one or more classes are significantly associated with multiple neuropsychiatric and neurodegenerative diseases. Most of these genes exhibit changes in expression profiles only within a single class, and nearly half of these genes show diverged expression limited to non-neuronal cell types. Since gene coexpression reflects shared regulation and function, we verify whether the observed gene expression changes have a functional impact by studying the divergence of gene coexpression relationships across species.

While single cell data captures the specificity of coexpression across individual cell types, bulk transcriptomic data is better powered to assess differential network connectivity across populations over a wider range of species that could explain the changes in gene activity observed at single cell resolution. We utilize species-specific gene coexpression networks to provide a quantitative framework to connect changes in gene expression profile to species-specific differential regulation. We identify 139 genes (<1% of all expressed genes) with human-specific expression and connectivity patterns not replicated in other primates or mammals. Relative to other expressed genes, these “human-divergent” genes are younger, and display significantly higher rates of sequence evolution and evolve under relaxed mutational constraint. We propose that integrating both types of data can detect both conserved genes that are well-suited for translational research, and genes with differential co-regulation across species that could limit their utility as disease biomarkers in model organisms (*66*). Further work is necessary to discern the molecular mechanisms underlying the human-specific changes in expression regulation of these genes. Additionally, we note that while our results are focused on genes with human-specific differential regulation, our datasets and framework can be extended to identify genes with differential regulation specific to other species or phylogenetic groups in general.

In summary, we generate a consensus taxonomy of cell types in the MTG of humans, two great apes, and two monkeys, to evaluate the conservation of gene function across primates. Further, we present a comprehensive catalog of coexpression conservation across metazoa which we combined with the gene expression data at single cell resolution to infer genes with changed activity in the human lineage. We infer a handful of genes with marked changes in both cell type-specific expression and gene coexpression neighborhoods, which could underlie evolutionary innovations exclusive to the human lineage. Overall, our datasets provide an opportunity to explore gene functional conservation at single cell resolution and across large evolutionary distances, and examine the regulatory divergence of genes associated with human-specific traits and diseases. We provide the datasets through a web-based tool (https://gillisweb.cshl.edu/Primate_MTG_coexp/) for users to explore and select genes with conserved or species-specific expression regulation.

## Methods

### Single nucleus RNA-sequencing processing and clustering

Over 570,000 nuclei were collected from five primates. All nuclei preparations were stained for the pan-neuronal marker NeuN and FACS-purified to enrich for neurons over non-neuronal cells. Samples containing 90% NeuN+ (neurons) and 10% NeuN- (non-neuronal cells) nuclei were used for library preparations and sequencing. Nuclei were included in downstream analysis if they passed all QC criteria:

SMART-seq v4 criteria:

> 30% cDNA longer than 400 base pairs
> 500,000 reads aligned to exonic or intronic sequence
> 40% of total reads aligned
> 50% unique reads
> 0.7 TA nucleotide ratio

Cv3 criteria:

> 500 (non-neuronal nuclei) or > 1000 (neuronal nuclei) genes detected
< 0.3 doublet score

Datasets from each species and modality (SSv4, Cv3) were analyzed independently. Briefly, each dataset was subset into 5 neighborhoods (CGE-derived and MGE-derived inhibitory neurons, IT type and deep excitatory neurons, and non-neurons) based on prior knowledge from human and mouse studies of cortical cell types (*2, 29*). Each neighborhood was annotated using the label transfer function from Seurat (*67*) with cell subclass labels from the recently published human primary motor cortex (M1) taxonomy (*29*). Subclass label transfer was performed using 3000 highly variable genes or their orthologs for human and non-human primate datasets, respectively. Datasets underwent additional QC and passing nuclei from each dataset were normalized using SCTransform (*68*). An integrated space was generated for each species by performing a canonical correlation analysis (CCA) across individuals and modalities. Each integrated space was clustered into hundreds of ‘metacells’, and metacells which passed quality control were merged with their nearest neighbors until merging criteria were met, resulting in the final clusters for each species (refer to https://github.com/AllenInstitute/Great_Ape_MTG for further details on RNA-seq processing, QC and annotation).

### Replicability of clusters

All analyses were performed in R version 4. MetaNeighbor v1.12 (*31, 32*) was used to provide a measure of neuronal and non-neuronal subclass and cluster replicability within and across species. We used OrthoDB v10.1 (*34*) to shortlist 14,131 orthologs across five primates, and subset snRNA-seq datasets from each species to this list of common orthologs before further analysis. For each assessment, we identified highly variable genes using the get_variable_genes function from MetaNeighbor. In order to identify homologous cell types, we used the MetaNeighborUS function with the fast_version and one_vs_best parameters set to TRUE. The one_vs_best parameter identifies highly specific cross-dataset matches by reporting the performance of the closest neighboring cell type over the second closest as a match for the training cell type, and the results are reported as the relative classification specificity (AUROC). This step identified highly replicable cell types within each species and across each species pair. All 24 subclasses are highly replicable within and across species (one_vs_best AUROC of 0.96 within species and 0.93 across species in **Fig. 1B**).

While cell type clusters are highly replicable within each species (one_vs_best AUROC of 0.93 for neurons and 0.87 for non-neurons), multiple transcriptionally similar clusters mapped to each other across each species pair (average cross-species one_vs_best AUROC of 0.76). To build a consensus cell type taxonomy across species, we defined a cross-species cluster as a group of clusters that are either reciprocal best hits or clusters with AUROC > 0.6 in the one_vs_best mode in at least one pair of species. This lower threshold (AUROC > 0.6) reflects the high level of difficulty/specificity of testing only against the best performing other cell type. We identified 86 cross-species clusters, each containing clusters from at least two primates. Any unmapped clusters were assigned to one of the 86 cross-species clusters based on their transcriptional similarity. For each unmapped cluster, top 10 of their closest neighbors are identified using MetaNeighborUS one_vs_all cluster replicability scores, and the unmapped cluster is assigned to the cross-species cluster in which a strict majority of its nearest neighbors belong. For clusters with no hits, this is repeated using top 20 closest neighbors, still requiring a strict majority to assign a cross-species type. A total of 594 clusters present in all five primates mapped to 86 cross-species clusters, with 493 clusters present in 57 consensus cross-species clusters shared by all five primates. Five of the 57 consensus cell types are visualized in the third panel in **Fig. 1C**. Gene expression across single nuclei present in 57 consensus clusters in the human MTG were visualized using UMAP plots colored by log-transformed expression levels (**Fig. 2C**).

### Calculating the expressolog score

We generated a pseudo-bulk dataset for each species which records the normalized average counts per cell type (CPM) of 14,131 genes across the consensus cell types. For each pair of species, we calculated the expression profile similarity for all pairs of genes by computing the Pearson correlation of normalized expression levels across 57 homologous cell types. For each gene in one species, we calculated the rank-standardized expression profile similarity of its 1:1 ortholog (relative to 14,130 genes) in the other species, repeated this calculation in the opposite direction, and report the average of the bidirectional scores as the “expressolog score” (see **Fig. 2B** for schematic representation). The expressolog score is equivalent to the average Area Under the Receiver-Operating characteristic Curve (AUROC), with a score of 1 indicating that orthologs can be identified by matching expression profiles across species, and a score of 0 suggesting that orthologs have diverged in expression across species. The AUROC for expression profile similarity of gene *i* in one species with gene *j* in another species is calculated as:

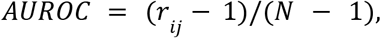

where *N* represents the total number of genes, and *r_ij_* is the rank of the Pearson correlation of gene *i* with gene *j* relative to other (*N - 1*) genes in the second species. In our expressolog analysis, *N* = 14,131.

Expressolog scores are also calculated across cell types within each class, subtype and meta-cluster (as defined in **Fig. 2A**) using the same formula in order to assess ortholog expression similarity at different levels of granularity. For each cell type group (ex: excitatory neurons, MGE-derived inhibitory neurons, or *Pvalb* meta-cluster), we obtain a 14,131 x 14,131 matrix of AUROCs corresponding to the expression profile similarity of all gene pairs across a pair of species, and report the AUROCs corresponding to 1:1 orthologs as expressolog scores. Average expressolog scores between human and non-human primates calculated within each cell type group are reported in **Table S3**. For each gene pair across a pair of species, overall expressolog scores at the class, subtype and meta-cluster levels capture the extent to which gene expression variation within relatively homogeneous cell types is shared across species, and are computed as follows. The 14,131 x 14,131 AUROC matrices for all cell type groups at a desired level of granularity are rank-standardized, aggregated and rank-standardized again, and the resulting values are reported as the overall expressolog scores for a gene pair at that level of granularity. Overall expressolog scores between human and non-human primates calculated at four levels of cellular heterogeneity are listed in **Table S4**.

### Isoform data generation

We used Smart-Seq v4 single nucleus RNA-seq data from human, chimp and gorilla to assess the expression profile similarity of individual isoforms across great apes. Reads from cells belonging to the consensus clusters were mapped to the species’ genomes using the default parameters in STAR v2.7.7a (*69*). Isoform and gene expression were quantified using RSEM v1.3.3. For the analysis related to Fig. 2, we retained consensus clusters with reads mapped from 10 or more cells, and further removed isoforms with total expression < 5 TPM. In order to assess whether an isoform could predict itself among other isoforms of a gene, we considered genes with at least two isoforms shared by all species. We computed the expressolog scores of all pairs of isoforms of a gene across a pair of species, and ranked the expressolog score of an isoform with itself relative to other isoforms (reported as an AUROC).

### Building aggregate coexpression networks

All coexpression networks used in this study were generated by aggregating networks built from individual cell types or datasets. In brief, networks for each cell type or dataset are built by calculating the Spearman correlation between all pairs of highly variable genes based on read counts, then ranking the correlation coefficients for all gene-gene pairs, with NAs assigned the median rank. Aggregate networks are generated by averaging rank standardized networks from individual datasets.

Single nucleus coexpression networks were generated by aggregating 57 cell type-specific networks. Meta-analytic coexpression networks derived by aggregating 54,668 individual RNA-seq datasets covering 21 metazoan species and yeast were downloaded from CoCoCoNet (*30*). Four RNA-seq datasets each were used to build aggregate coexpression networks for gorilla and marmoset. Human bulk brain coexpression network was generated by aggregating 20 individual datasets curated by GEMMA (*46*). To assess the connectivity of marker gene groups in coexpression networks, we used the *run_neighbor_voting* function from the EGAD R package (*44*).

### Curated gene sets and orthology

To investigate the conservation and divergence of the coexpression of gene families between human and non-human primates, we carried out MetaNeighbor analysis using gene groups curated by the HUGO Gene Nomenclature Committee (HGNC) at the European Bioinformatics Institute (https://www.genenames.org; downloaded October 2021) and by the Synaptic Gene Ontology (SynGO (*70*), downloaded October 2021). HGNC annotations were propagated via the provided group hierarchy to ensure the comprehensiveness of parent annotations. Only groups containing five or more genes were included in the analysis.

The MetaMarkers package (*71*) was used to find marker genes for cell types defined at different levels of organization in each species, with search at each level stratified by the broader cell type so as to generate marker sets that can discriminate even relatively homogeneous cell clusters. Marker genes defining cell class, subclass and consensus clusters are listed in **Table S5**. List of transcription factors used in Fig. 2 were obtained from Ziffra et al (*62*).

To assess genic features associated with human-divergent genes, we downloaded the sequence similarity, gene length and GC content for all human genes from Ensembl v107, gene ages from GenTree (http://gentree.ioz.ac.cn/), list of significant eQTLs and associated genes from the GTEx portal (v8, (*54*)), and gene constraint scores (LOEUF) from The Genome Aggregation Database (gnomAD v2.1.1). Average sequence evolution rates between human and other primates, and gene lists associated with various brain disorders were downloaded from the GenEvo website (https://genevo.pasteur.fr/). Gene set enrichment analysis was performed using Fisher’s exact test and the resulting p-values were adjusted by applying Benjamini-Hochberg correction. Cell type-specificity scores were calculated using pseudo-bulk human MTG data as published (*72*).

OrthoDB v10.1 (*34*) was used for orthology mapping. For each pair of species, we used the set of orthology groups of their last common ancestor to obtain a comprehensive list of many-to-many orthologs. We filtered this list to include only 1:1 orthologs, which yielded ~ 4500 orthologs for phylogenetically distant species (like human and yeast) and ~ 13,500 orthologs for recently diverged species. All single nucleus expression profile similarity analyses used a set of 14,131 orthologs across five primates, with aggregate coexpression networks built using a subset of the top 4,500 highly variable genes. Species divergence times were sourced from TimeTree (*73*).

### Calculating cross-species coexpression conservation

For each pair of species to be compared, we filter aggregate coexpression networks to include known 1-1 orthologous genes, then we compare each gene’s top 10 coexpression partners across species to quantify gene functional similarity (*43*). The schematic in **Fig. 3B** illustrates the calculation of coexpression conservation of an orthologous gene between human and rhesus macaque. Given a gene of interest in human (yellow circle on the left), its top 10 coexpression partners are identified and coexpression conservation is calculated by ranking the coexpression of their macaque orthologs with the macaque ortholog of the target gene (yellow circle on the right). Calculation is repeated in the other direction (macaque to human), and the average of bi-directional AUROC is taken as a measure of coexpression neighborhood similarity of the target gene. We calculate the coexpression conservation not just for orthologs, but for all gene pairs, and rank the coexpression conservation of each ortholog relative to all genes to determine the specificity of coexpression neighborhood conservation for each gene. We term this specificity score as “coexpression conservation” and note that it provides a standardized measure to compare the extent of functional conservation of orthologs over large evolutionary timescales, and infer examples of human-specific regulatory divergence.

### Protein sequence similarity for candidate genes showing regulatory divergence in humans

We obtained data for protein sequence similarity of 1:1 orthologs between humans and non-human primates from Ensembl v107. *NHEJ1* - 96% and *GTF2H2* - 91% average similarity between human and great apes (chimp, gorilla) and monkeys (crab-eating macaque, rhesus macaque). *C2* - 97.87% similarity of humans with crab-eating macaque. *BBS5* - 99.7% similarity between human and two great apes.

## Acknowledgements

We thank Leon French, Stephan Fischer, Risa Kawaguchi, Michael Passalacqua and John Hover for thoughtful feedback on the manuscript.

## Funding

National Institutes of Health grant R01LM012736 (HS, JG)

National Institutes of Health grant R01MH113005 (HS, JG)

National Institutes of Health grant U19MH114821 (HS, JG)

National Institutes of Health grant F32MH114501 (MC)

NARSAD Young Investigator Award (MC)

National Institutes of Health grant R01HG009318 (AD)

National Institutes of Health grant U01MH114812 (NJ, RH, EL, TB)

## Author contributions

Sample preparation and RNA data generation: NJ, RH, EL, TB

Isoform data generation: AD

Conceptualization: JG

Data archive / Infrastructure: HS

Data analysis: HS

Data interpretation: HS, MC, AD, JG

Writing manuscript: HS, MC, JG

## Competing interests

Authors declare that they have no competing interests.

## Data and materials availability

Integrated snRNA-seq gene expression dataset and associated metadata for each primate species are available on the BICCN Human/NHP website. Data and code to reproduce the cross-species expression profile similarity and coexpression conservation analysis can be accessed from https://labshare.cshl.edu/shares/gillislab/resource/Primate_MTG_coexp/ and https://github.com/hamsinisuresh/Primate-MTG-coexpression, respectively.

An online tool that allows researchers to explore (i) the expressolog score of orthologs between human and non-human primates at different cell type resolutions, and (ii) the functional conservation of orthologs across metazoan species is available through our web server https://gillisweb.cshl.edu/Primate_MTG_coexp/. Classification of 14,131 genes based on their coexpression divergence in single cell, bulk or both transcriptomic datasets is provided in **Table S7**.

## Supplementary Materials

Figs. S1 to S5

Tables S1 to S7

## Notes

https://gillisweb.cshl.edu/Primate_MTG_coexp/

https://github.com/gillislab/Primate-MTG-coexpression

### Competing Interest Statement

The authors have declared no competing interest.

