## Supplementary Figures S1 to S5 for "Conserved coexpression at single cell resolution across primate brains"

**This PDF file includes:**

Figs. S1 to S5

Tables S1 to S7

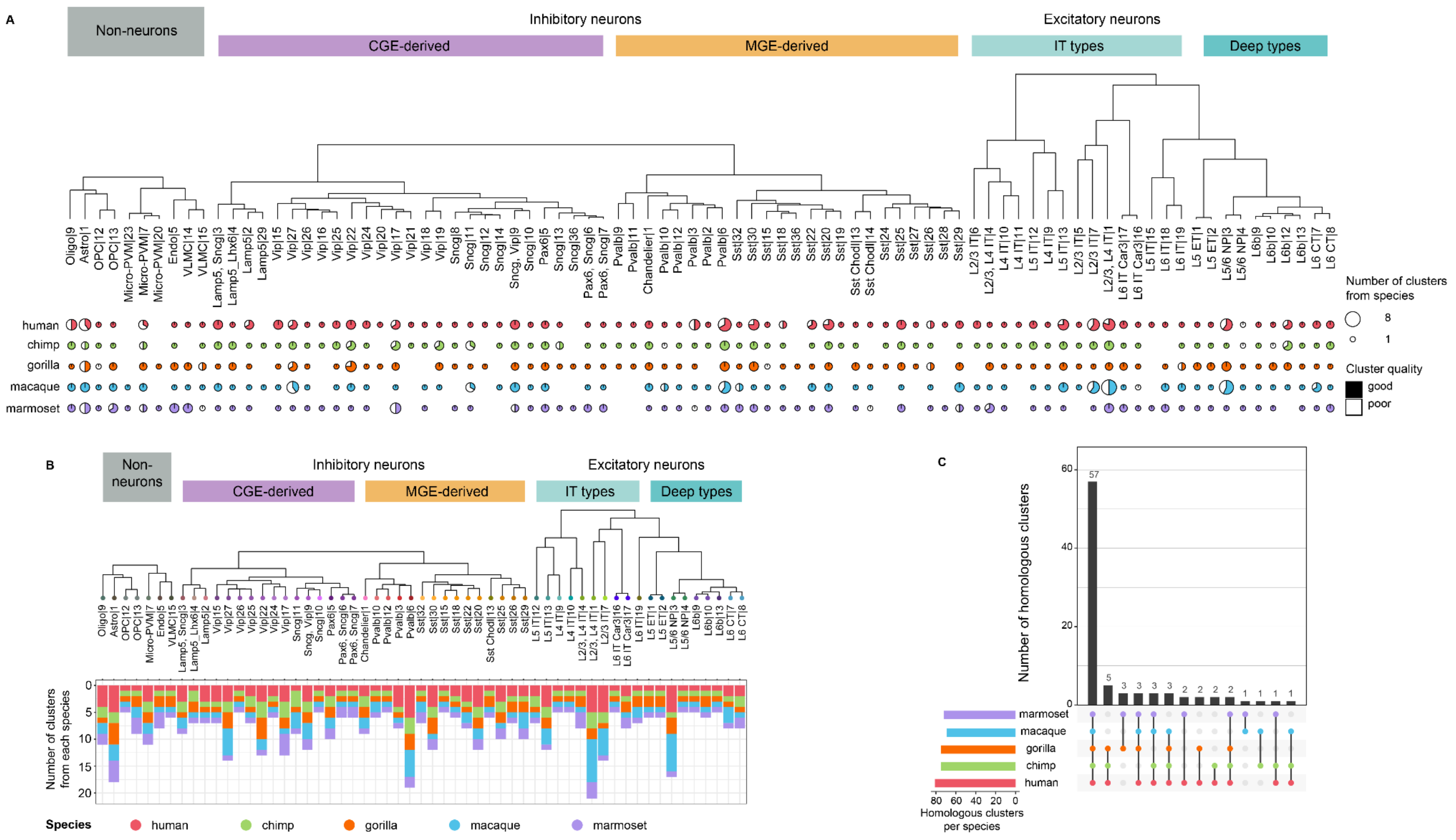


**Fig. S1. Constructing consensus taxonomy of homologous cell types.** (**A**) Complete cross-species cluster dendrogram generated using average-linkage hierarchical clustering with (1 - average MetaNeighborUS one_vs_all cluster replicability scores) for each pair of 86 cross-species clusters as a measure of distance between cell types. (**B**) Consensus cross-species taxonomy is visualized by pruning the dendrogram (shown in (**A**)) to retain only the 57 homologous cell types shared by all five species. (**C**) Upset plot showing the distribution of 86 homologous cell types across species.

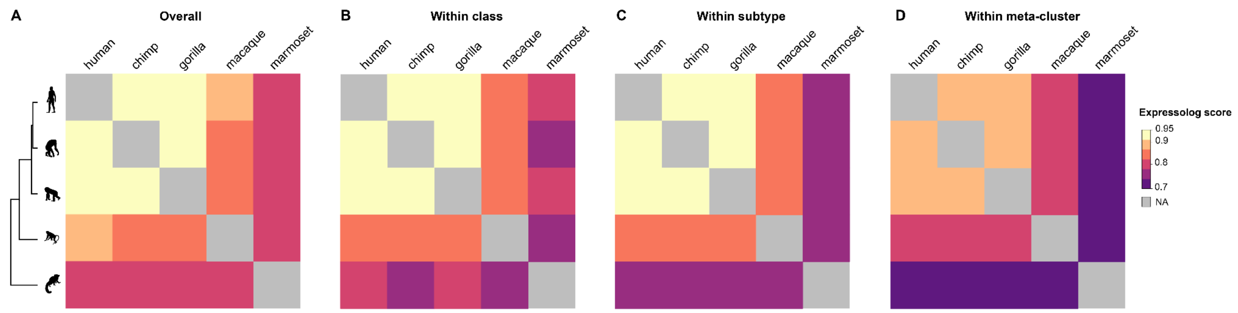


**Fig. S2. Expression profile similarity correlates with species phylogeny even when stratified by cell-type hierarchy.** Heatmaps indicate the mean expression profile similarity of orthologs between each species pair calculated across (**A**) 57 consensus cell types, (**B**) cell types within each class, (**C**) cell types within each subtype, and (**D**) cell types within each meta-cluster. List of cell types at different resolutions are provided in **Fig. 2A**.

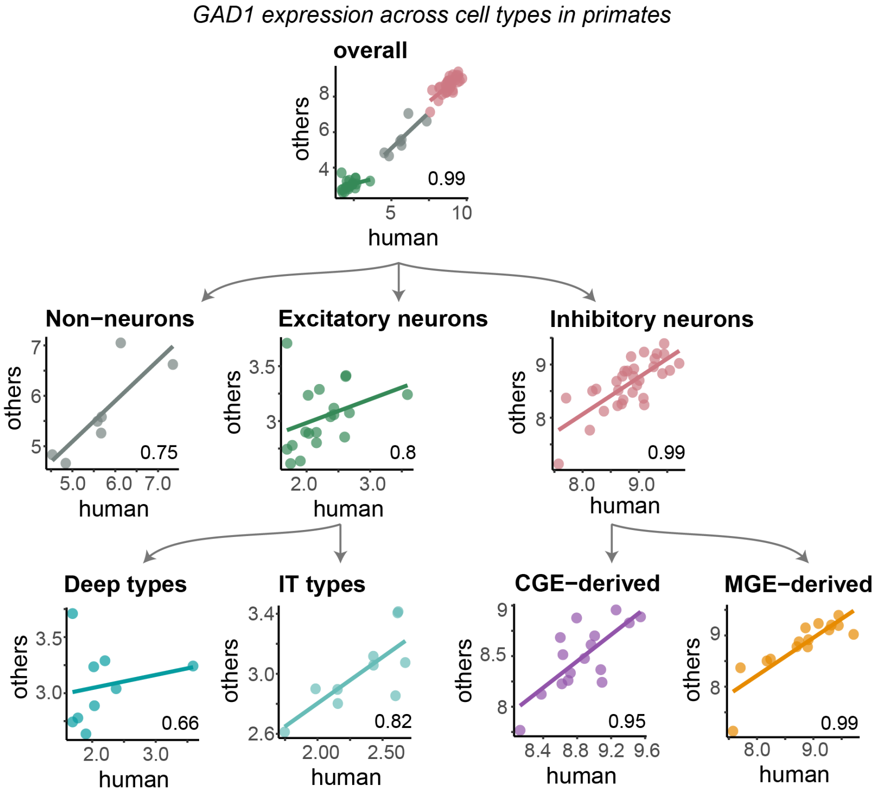


**Fig. S3. Inhibitory neuron marker GAD1 shows continuous variation in expression across cell-types in human and non-human primates.** Scatter plots show the expression level (log2 CPM) of GAD1 in human and non-human primates across cell types at different resolutions (expressolog score at bottom right).

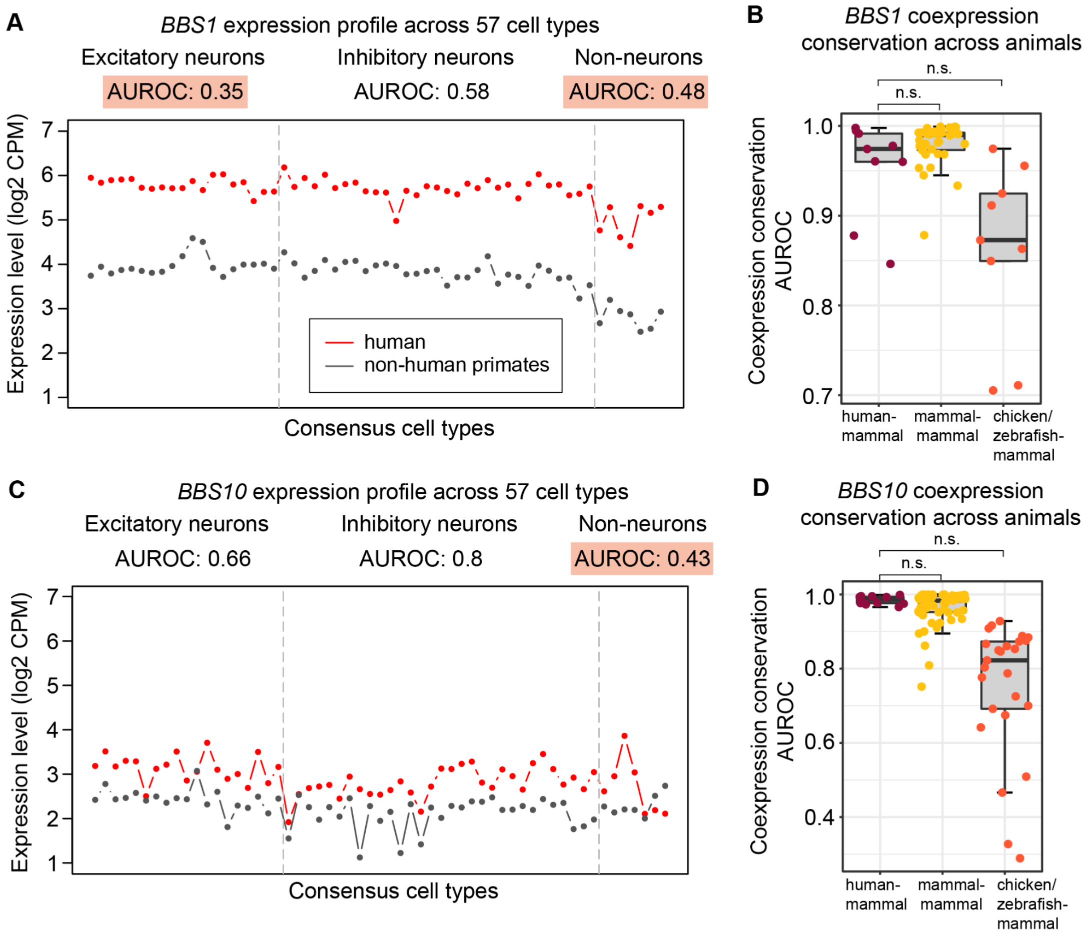


**Fig. S4. BBS1 and BBS10 show diverged expression in one or more classes in humans, but have conserved coexpression neighborhoods across metazoa.** (**A, C**) Plots compare the expression profile of each gene of interest in humans with the average expression profile of the ortholog in non-human primates. Expressolog scores within each cell class are listed above the plot, and scores < 0.55 are highlighted in orange. (**B, D**) Boxplots show coexpression conservation for orthologs between human and non-human mammals (points colored in maroon), between pairs of non-human mammals (yellow), and between non-human mammals and other vertebrates (chicken and zebrafish; orange).

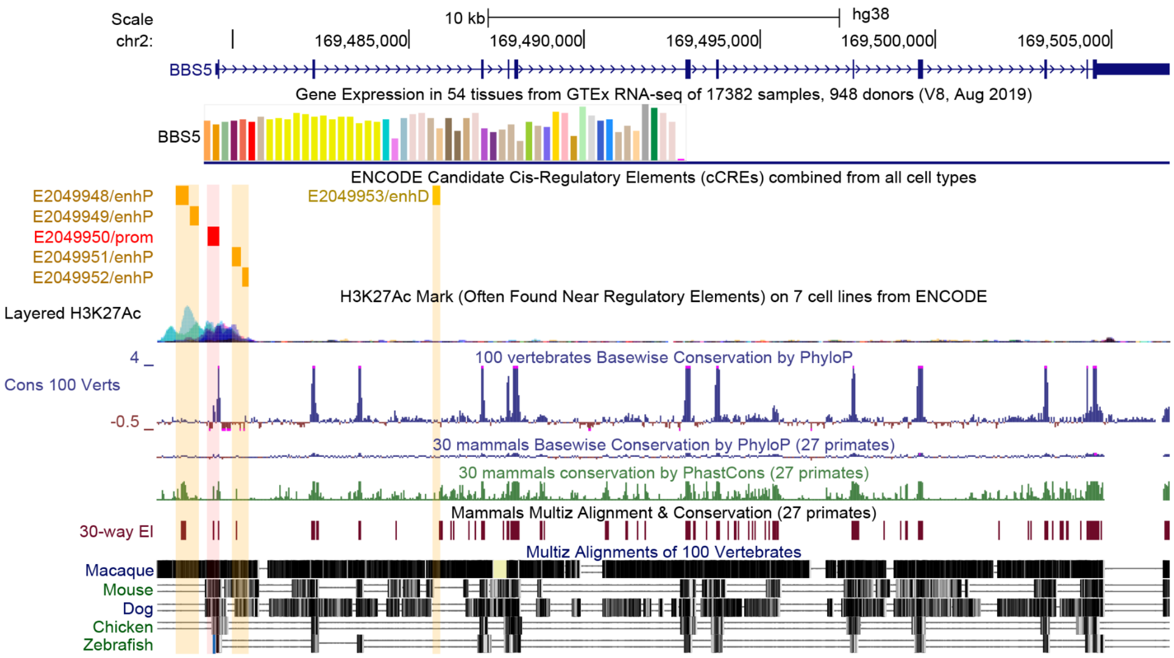


**Fig. S5. BBS5 candidate CREs from ENCODE.** High MULTIZ alignment score with macaque and low vertebrate PhyloP scores indicate that putative enhancers regulating human BBS5 expression are conserved within primates, but diverged across mammals and other vertebrates.

### Supplementary Tables

**Table S1: Summary of single-nucleus transcriptomic datasets from the middle temporal gyrus (MTG) of five primates**

Summary of metadata (i.e. number of nuclei, number of donors, sequencing technology and number of within-species clusters) associated with the single-nucleus transcriptomic datasets generated from the middle temporal gyrus from five primates.

**Table S2: Performance of 920 HGNC and SynGO gene sets in classifying consensus cell types within and across primates**

Classification scores are calculated for gene sets containing 5 or more genes, in each cell type within and across primates. The scores are averaged over species and cell types are reported as 'within_species' and 'cross_species' mean AUROC.

**Table S3: List of expressolog score calculated across cell types at different levels of cell hierarchy**

Average expressolog scores calculated between human and non-human primates across cell types within each class, subtype and meta-cluster.

**Table S4: List of expressolog scores (expression profile similarity of orthologs relative to other genes) at different levels of cellular organization**

Expressolog scores (AUROC) for 14,131 genes are calculated at different levels of cell type hierarchy to measure the extent of conserved cross-species coexpression at multiple scales. Average scores computed over all primate pairs are reported in the first four columns and average scores between human and non-human primates in the last four columns.

**Table S5: List of marker genes for cell classes, subclasses and cell types in the human MTG transcriptomic data**

Marker genes for 3 cell classes, 24 subclasses and 57 cell types were selected using the MetaMarkers package. We identified 200, 100, 50 genes for each cell class, subclass and cell type respectively, indicated by '1' in the table.

**Table S6: Metadata related to cross-species coexpression conservation calculation**

List of species and metadata related to their aggregate coexpression networks used to calculate coexpression conservation of 14,131 genes across animal kingdom.

**Table S7: 14,131 genes classified into different categories based on their potential for human-specific functional divergence identified using single-cell and bulk transcriptomic data**

Genes are classified as:

I. 'Diverged_in_single-cell' if their average expressolog score between human and non-human primates drops below 0.55 in one or more classes

II. 'Diverged_in_bulk' if the distribution of gene coexpression conservation between human and non-human mammals is significantly lower than that within mammals, and between vertebrates (chicken and zebrafish) and non-human mammals

III. 'Diverged_in_both' for 139 genes with concordant divergence in single-cell and bulk data (i.e. I and II are TRUE)

IV. NA, otherwise
